# Characteristic sounds facilitate object search in real-life scenes

**DOI:** 10.1101/563080

**Authors:** Daria Kvasova, Laia Garcia-Vernet, Salvador Soto-Faraco

## Abstract

Real-world multisensory events do not only provide temporally and spatially correlated information, but also semantic correspondences about object identity. Semantically consistent sounds can enhance visual detection, identification and search performance, but these effects are always demonstrated in simple and stereotyped displays that lack ecological validity. In order to address identity-based crossmodal relationships in real world scenarios, we designed a visual search task using complex, dynamic scenes. Participants searched objects in video clips from real life scenes with background sounds. Auditory cues embedded in the background sounds could be target-consistent, distracter-consistent, neutral and no sound (just background noise). We found that characteristic sounds enhance visual search of relevant objects in natural scenes but fail to increase the salience of irrelevant distracters. Our findings generalize previous results on object-based crossmodal interactions with simple stimuli and shed light upon how audio-visual semantically congruent relationships play out in real life contexts.

## 1. Introduction

Interactions between sensory modalities are at the core of human perception and behavior. For instance, the distribution of attention in space is guided by information from different sensory modalities as shown by cross-modal and multisensory cueing studies (e.g., Spence & Driver, 2004). Most research on crossmodal interactions in attention orienting has typically employed the manipulation of spatial (Spence & Driver, 1994; Driver & Spence, 1998; McDonald, Teder-Salejarvi, & Hillyard, 2000) and temporal (Busse et al., 2005; Van der Burg, Olivers, Bronkhorst, & Theeuwes, 2008; Van den Brink, Cohen, van der Burg, Talsma, Vissers, & Slagter, 2014; Maddox, Atilgan, Bizley, & Lee, 2015) congruence between stimuli across modalities. However, recent studies have highlighted that in real world scenarios, multisensory inputs do not only convey temporal and spatial congruence, but also bear semantic relationships. The findings of these studies have shown that crossmodal correspondences at the semantic level can affect detection and recognition performance in a variety of tasks, including the distribution of spatial attention (e.g., Chen & Spence, 2011; Molholm, Ritter, Javitt, & Foxe, 2004; Pesquita, Brennan, Enns, & Soto-Faraco, 2013; Iordanescu, Guzman-Martinez, Grabowecky, & Suzuki, 2008; Iordanescu, Grabowecky, Franconeri, Theeuwes, & Suzuki, 2010; List, Iordanescu, Grabowecky, & Suzuki, 2014). For instance, in visual search amongst images of everyday life objects, sounds that are semantically consistent (albeit spatially uninformative) with the target speed up search times, in comparison to inconsistent or neutral sounds (Iordanescu, et al., 2008; Iordanescu, et al., 2010). However, one paramount question which remains to be answered in this field is, to which extent such multisensory interactions discovered under simplified, laboratory conditions, have an impact under the complexity of realistic, multisensory scenarios (Matusz et al. 2019; Soto-Faraco et al., in press). We set out to address this question.

Previous findings on crossmodal semantic effects on search behaviour so far have used static, stereotyped artificial scenarios that lack meaningful context (Iordanescu, Guzman-Martinez, Grabowecky, & Suzuki, 2008; Iordanescu, Grabowecky, Franconeri, Theeuwes, & Suzuki, 2010; List, Iordanescu, Grabowecky, & Suzuki, 2014). However, searching targets in these simplified displays used in laboratory tasks is very different from the act of looking for an object in complex, naturalistic scenes. As many authors have pointed out before, generalization of laboratory findings using idealized materials and tasks is often far from trivial (Matusz et al., 2019, for a recent review). Outcomes that are solid and replicable under these simplified conditions may turn out differently in contexts that are more representative of real-life (Maguire, 2012; Wolfe et al., 2005; Peelen & Kastner, 2014; for examples in visual research; see Soto-Faraco et al., in press, for a review concerning multisensory research). First, realistic scenes are usually far more cluttered than stereotyped search arrays. Second, natural scenarios provide organization based on relevant prior experience: When searching for your cat in the living room, you would not expect the cat hovering midway to the ceiling, next to a floating grand piano. Yet, many laboratory tasks require just that: A picture of a (target) cat can be presented within a set of randomly chosen objects that have no relations between them, arranged in a circle, against a solid white background (see Figure 1).

**Figure 1.**
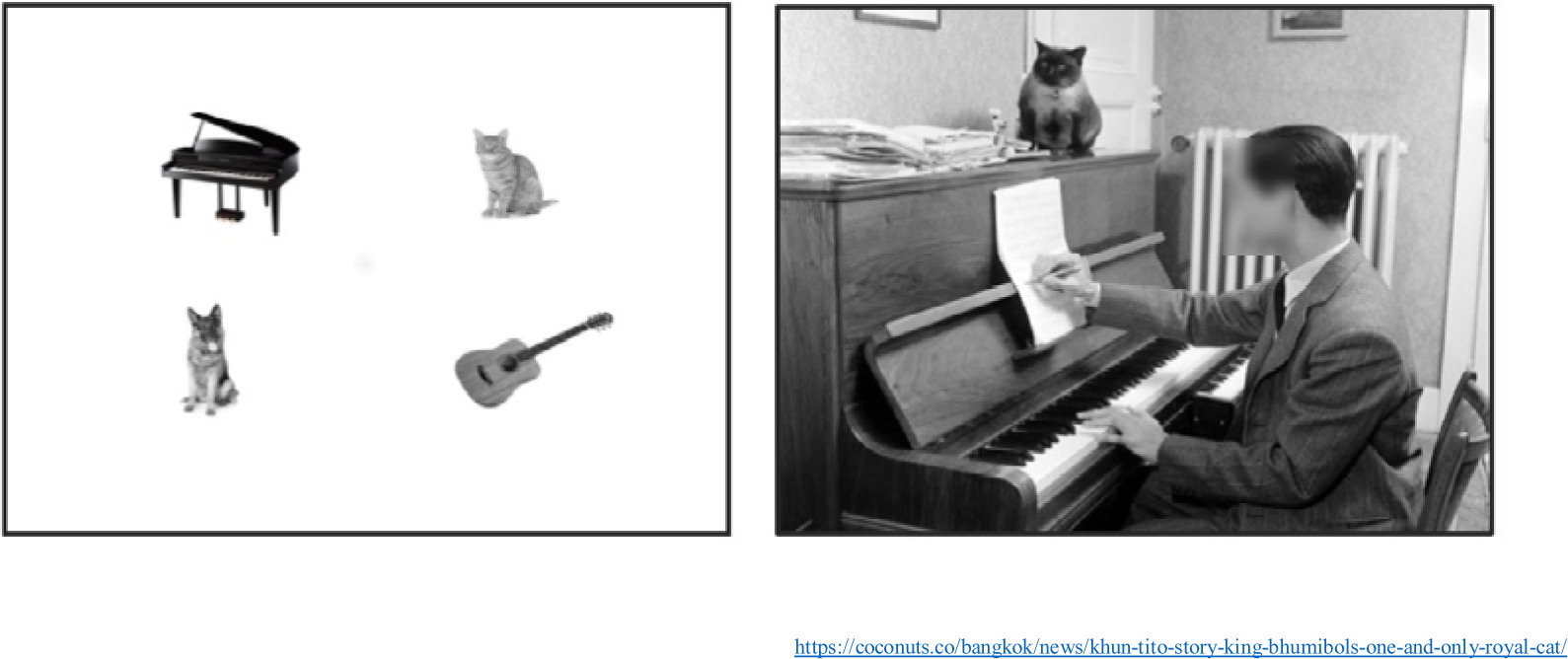
Left picture is an example of stimuli used as a typical search array in a search experiment. Figures are randomly chosen and randomly distributed in space without any meaningful connection between them. On the right naturalistic picture some objects are the same as on the left but now they are put into a context with spatial envelope, proportionality and variety of meaningful and functional connections between objects

Previous visual-only studies have already made a point about the differences in how spatial attention is distributed in naturalistic, real life scenes compared to simple artificial search displays typically used in psychophysical studies (e.g., Peelen & Kastner, 2014, for a review; Henderson & Hayes, 2017). Given that experience and repetition tends to facilitate visual search (Shiffrin & Schneider, 1977; Evans, Georgian-Smith, Tambouret, Birdwell, & Wolfe, 2013; Kuai, Levi, & Kourtzi, 2013), another important difference could lie in our familiarity (and hence, predictability) with natural scenes, compared to laboratory displays. In addition, humans can extract abundant information from natural scenes (gist) at a glance, quickly building up expectations about the spatial layout and relationships between objects (Biederman, Mezzanotte, & Rabinowitz, 1982; Greene & Oliva, 2009; Peelen, Fei-Fei, & Kastner, 2009; MacEvoy & Epstein, 2011).

For example, Nardo, Santangelo & Macaluso (2014) reported that crossmodal semantic congruency between visual events and sounds had no effect on spatial orienting or brain activity during free viewing of videos from everyday life scenes. In contrast, another study by Mastroberardino, Santangelo & Macaluso (2015) with static images reported that visual images could capture spatial attention when a semantically congruent, albeit spatially uninformative sound was presented concurrently. Along a similar line, Iordanescu et al. (2008; 2010) showed that spatially uninformative characteristic sounds, speeded up visual search when consistent with the visual target. Conversely to the study of Nardo et al (2014), which found no effect, Iordanescu et al. (2008; 2010) and Mastroberardino et al. (2015) used simple static images presented in decontextualized search arrays (Iordanescu, et al., 2008; Iordanescu et al., 2010). Both, these differential features (dynamic nature of natural scenes and their complexity) have been pointed out as important components for the generalization of cognitive psychology and neuroimaging findings to real-world contexts (e.g., Hasson, Malach, & Heeger, 2010). Another possibly important variable in prior research on cross-modal semantic influence on attention is task-relevance. Unlike Nardo et al. (2014) and Mastroberardino et al. (2015) studies, in the study of Iordanescu et al. (2008, 2010) the critical (target) objects were task-relevant, potentially making audio-visual congruence relations also relevant to the task.

Based on the results of these prior studies, one first outstanding question is whether crossmodal semantic relationships can play a role at all in complex dynamic scenarios. Until now, the only study using such scenarios (Nardo et al., 2014) has returned negative results, in contrast with other studies using more stereotypical displays (Iordanescu et al., 2008, 2010; Mastroberardino et al., 2015). Given that a major difference between these studies was task relevance of the cross-modal events, a second interrelated question is whether the impact of crossmodal semantic relationships, if any, is limited to behaviourally relevant events. Here we present a study using a novel search task on realistic scenes, in order to shed light on these two questions.

In our visual search protocol, targets were everyday life objects appearing in video clips of naturalistic scenes. Spatially-uninformative characteristic sounds of objects mixed with ambient noise were presented during search. The relationship between the object sounds and the visual target defined four different conditions: *target-consistent sound, distracter-consistent sound, neutral sound* and *no sound*, which was a baseline condition that contained only background ambient noises. Visual search performance was measured with reaction times.

We hypothesised that, if crossmodal semantic congruency guides attention in complex, dynamic scenes, then reaction times should be faster in the target-consistent condition than in the distracter-consistent, neutral or no sound conditions (e.g., target-consistent characteristic sounds will help attract attention to the corresponding visual object). Regarding the possible task-relevance modulation of crossmodal semantic effects, we hypothesized that if audio-visual semantic congruence attracts attention in natural scenes automatically even when the objects are irrelevant to the current behavioural goal, then one should expect a slowdown in responses to targets in distracter-consistent trials, with respect to neutral sound trials. Else, if audio-visual semantic congruence has an impact only when task relevant (as we expected), then distractor-congruent sounds should not slow down performance compared to other unrelated sounds. In order to check the potential unspecific effects of object sounds on visual search times, such as alerting (Nickerson, 1973), we included neutral sound condition as a control. Neutral sounds were sounds that did not correspond to any object in the video of the current trial. Thus, we expected that differences due to general alerting of sounds, if any, would equally affect target-consistent, distractor-consistent, and neutral sound conditions, but not the no-sound baseline.

## 2. Method

### 2.1. Participants

Thirty-eight volunteers (12 males; mean age 25.22 years, SD = 3.97) took part in the study. They had normal or corrected-to-normal vision, reported normal hearing and were naïve about the purpose of the experiment. All subjects gave written informed consent to participate in the experiment. Two subject-wise exclusion criteria were applied before any data analysis. (1) If false alarm rate in catch trials (trials in which the search target was not present) was above 15%. (2) If accuracy in one or more conditions was below 70%. After applying these criteria, we retained data from thirty-two participants.

### 2.2. Stimuli

#### 2.2.1. Visual stimuli

A set of 168 different video-clips were obtained from movies, TV shows and advertisement, and others were recorded by experimenters from everyday life scenes. The video clips, size 1024×768 pixels and 30 fps were edited with Camtasia 9 software (https://www.techsmith.com/camtasia/) to 2 s duration fragments. No fades were used during the presentation. Ninety-six videos were used for the experimental conditions described below, and 72 videos for catch trials. For all of the videos, the original soundtrack was replaced with background noise created by superposition of various everyday life sounds (see example video clips and sounds in online supplemental materials).

Each video clip used for experimental (target-present) conditions contained two possible visual targets, which were always visual objects which have a characteristic sound (such as musical instruments, animals, tools, etc.). The criteria to choose the target objects in the videos was that, although they were visible (no occlusions, good contrast), they were not part of the main action in the scene. For instance, if a person is playing a guitar and this is the main action of the scene, the guitar could not be a target object. However, in a scenario with a band playing different instruments, the guitar could be a possible target. We applied this criterion to make the search non-trivial. Nevertheless, in order to compensate for potential biases related to particular objects or videos, we counterbalanced the materials so that each video and object contributed as target and as distractor in equal proportions across participants (see 2.3. Procedure).

#### 2.2.2. Auditory stimuli

We used characteristic sounds that corresponded semantically to the target/distractor objects (e.g. barking dog). However, they gave no information about the location of the object (sounds were always central) or its temporal profile (the sound temporal profile did not correlate with visual object motion or appearance). All the sounds were normalized to 89 dB SPL and had duration of 600 msec. Sounds were delivered through two loudspeakers placed at each side of the monitor, in order to render them perceptually central.

### 2.3. Procedure

The experiment was programmed and conducted using the Psychopy package 1.84.2 (Python 2.7) running under Windows 7. Participants were sitting in front of a computer monitor 22.5’’ (Sony GDM-FW900) at a distance of 77cm. We calibrated the video and sound onset latencies using The Black Box Toolkit (http://www.blackboxtoolkit.com, UK), within an error of SD=7.34ms.

In order to start each block of the experiment, participants pressed the space bar. Each trial started with a cue word printed on the screen indicating the target of the visual search for that trial. After 2000 ms, a video clip with the background noise plus, if applicable, a characteristic object sound of the corresponding condition (target-consistent, distracter-consistent, neutral) were presented. Following previous laboratory studies that used complex sounds and visual events we decided to desynchronize presentation of audiovisual event, by presenting the sound 100 ms before the video onset (Vatakis & Spence, 2010, for review; Knoeferle K. M., Knoeferle P., Velasco, & Spence, 2016, for a similar procedure).

The participant’s task was to judge whether or not the pre-specified target object was present in the video clip as fast as possible and regardless of its location. If the video ended before participants’ response, a question mark showed up on the screen and stayed there until participant responded. The next trial started 200 ms after the participant had responded [Fig. 2]. Half of the participants had to press A key (QWERTY keyboard) as soon as they found the target object. In case the object was not present on the scene, they pressed L key. For the other half it was the other way around. Visual search performance for each subject and condition was determined by the mean response time (RT) of correct responses.

**Figure 2.**
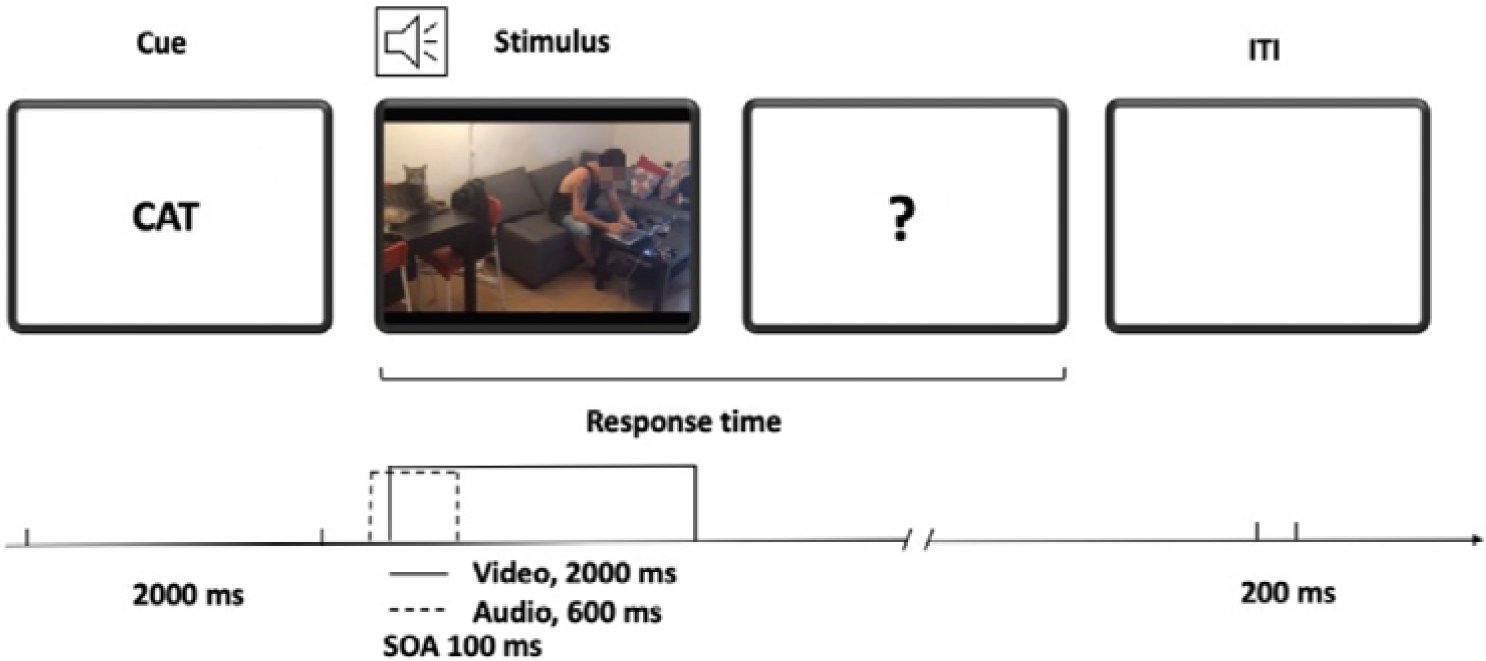
Sequence of events in the experiment. The trial started with the presentation of target word for 2000 ms. The target word was followed by the auditory cue and video. Auditory cue was presented 100 ms before the video was shown (SOA = 100ms) and lasted for 600ms while the video lasted for 2000 ms. There was no time limit for the participant response. 200ms after the participant had responded a new target word was presented.

Four types of sound-target conditions were used: *target-consistent, distractor-consistent, neutral* and *no sound*. In the target-consistent condition the identity of the sound matched with the target object. In the distractor-consistent condition the sound matched a non-target (distracter) object present in the scene. In the neutral condition the object sound did not match any of the objects in the scene. Finally, in the baseline condition, no particular object sound (auditory cue) was present (besides the background noise) [Fig. 3].

**Figure 3.**
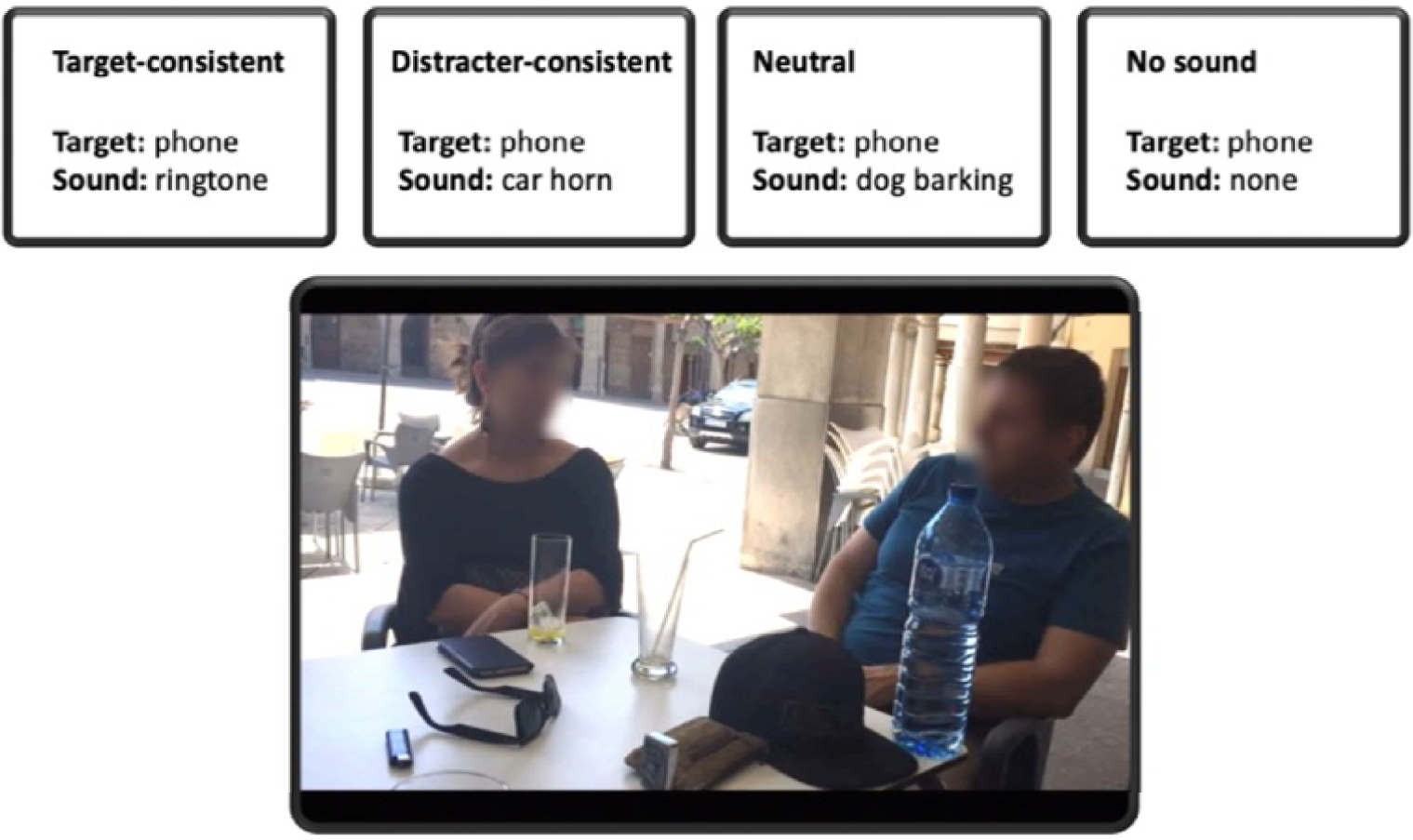
Example of conditions. In this example of stimulus, the possible targets are a mobile phone and a car. If the target is a mobile phone, in the target-consistent condition the sound will match the target, in the distractor-consistent condition the sound will match the distractor (a car), in the neutral condition the sound will not match any object of the scene (e.g. dog

Each participant saw each video-clip once, but overall, each video clip appeared in each of the four experimental conditions the same number of times (across subjects), except for catch trials which were the same for all participants. To achieve this, we created a total of 8 different versions of the experiment (in order to equate the number of times each of the two objects in each video was the target). In order to make sure that participants understood the task, they ran a 14-trial training block before the beginning of the experiment. The training set used video clips that were equivalent to, but not contained in, the experiment and included examples of the four experimental conditions as well as catch trials.

The experiment contained a total of 168 trials (24 trials per experimental condition plus 72 catch trials; hence the overall proportion of target present trials was ∼57%). The experiment was divided in six blocks of 28 videos with a representative number of trials of each condition and catch. Each participant received a different random order of videos.

## 3. Results and discussion

We ran a repeated measures ANOVA on mean RTs (for correct responses), with subject as the random effect and condition as the factor of interest. The analysis returned a significant main effect of condition (F(3,93)=3.14; p=0.0289). We further used paired t-tests to follow up on this main effect. Participants were faster in the target-consistent condition in comparison to distracter-consistent condition [t(31)=2.36, p<0.05, Cohen’s d=0.27], neutral [t(31)=2.33, p<0.05, Cohen’s d=0.39] and no sound [t(31)=2.53, p<0.01, Cohen’s d=0.32]. No difference was found between distracter-consistent, neutral and no sound conditions (see Figure 4). To ensure that there was no speed-accuracy trade off we analysed error data. The analysis showed that there was no difference in performance between conditions (Figure 5).

**Figure 4.**
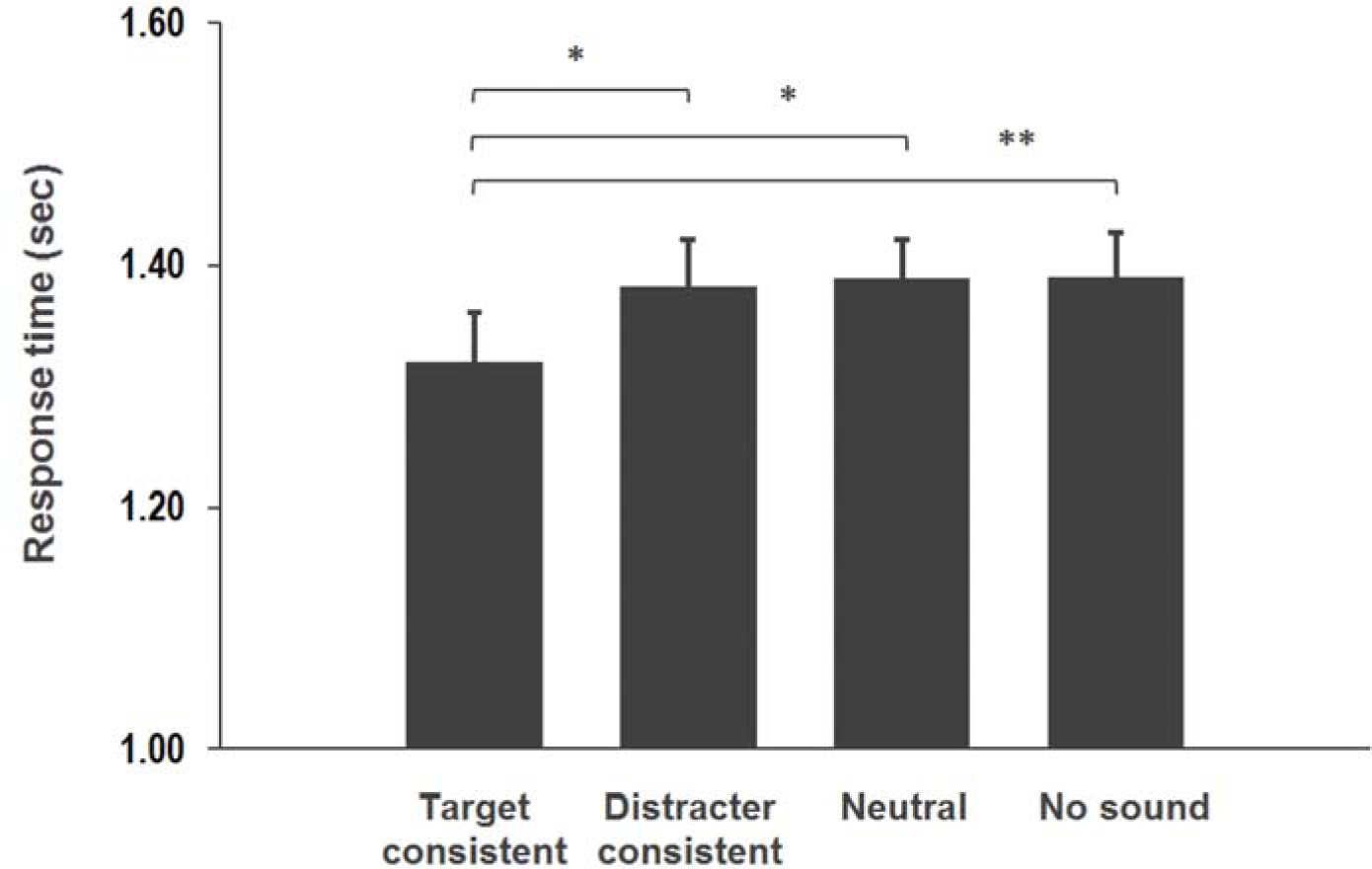
Visual search reaction times towards a target and error rates were plotted in the *target-consistent* sounds (black), *distracter-consistent* sounds (gradient), *neutral* sounds (grey) and *no sound* (white) conditions. Error bars indicate the standard error. Asterisks indicate significant difference between conditions (1 asterisk for p-value less than 0.05, 2 asterisks for p-value less than 0.01)

**Figure 5.**
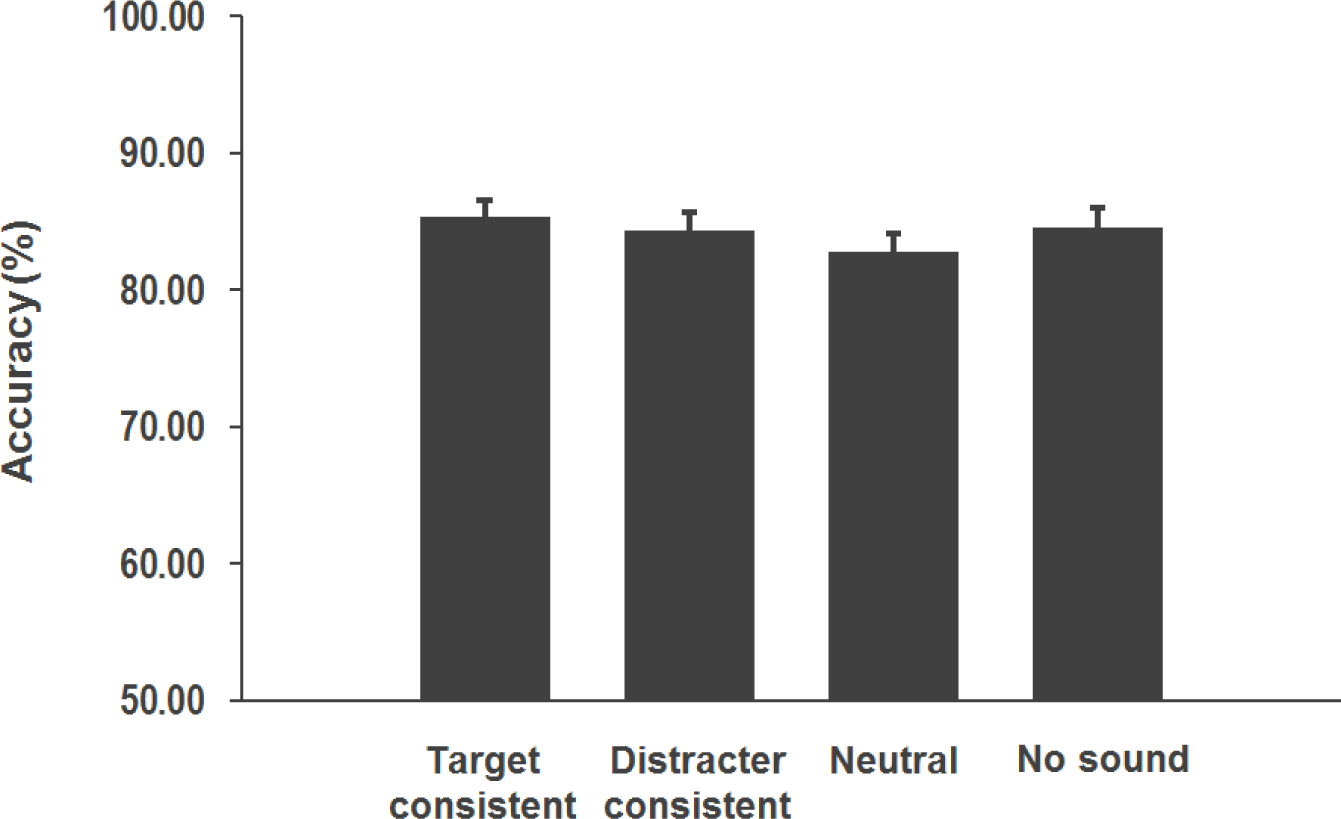
Visual search accuracy towards a target and error rates were plotted in the *target-consistent* sounds (black), *distracter-consistent* sounds (gradient), *neutral* sounds (grey) and *no sound* (white) conditions. Error bars indicate the standard error.

The results to emerge from the present study show that, when searching for objects in real life scenes, target-consistent sounds speed up search latencies in comparison to neutral sounds or when only background noises are present. Instead, distracter-consistent sounds produced no measurable advantage or disadvantage with respect to these baseline conditions (albeit, responses were slower than for target-consistent conditions). This finding demonstrates, for the first time, that characteristic sounds improve visual search not only in simple artificial displays (Iordanescu, et al., 2008; 2010) but also in complex dynamic visual scenes with contextual information. In general, and according to previous studies (Iordanescu, et al., 2008; Iordanescu et al., 2010), we can affirm that the results obtained in this study are due to object-based and not due to spatiotemporal correspondences, since we avoided any kind of spatiotemporal congruence. Semantic relationships between the objects in a complex visual scene can guide attention effectively (Wu, Wick, & Pomplun et al., 2014, for review), our results suggest that this semantic information did not make congruent auditory information redundant. Semantically consistent sounds can indeed benefit visual search along with available visual semantic information. This is the novel contribution of this study.

Despite research on attention orienting has been dominated primarily by low-level spatial and temporal factors (salience), recent research has focused on the role of higher-level, semantic aspects (e.g., Henderson & Hayes, 2017). Visual-only studies have highlighted, for example, the importance of functional relationships between objects (Biederman et al., 1982; Oliva & Torralba, 2007; MacEvoy & Epstein, 2011), expectancies regarding frequent spatial relations (Peelen et al. 2014, for review), and cues to interpersonal interactions (Kingstone, Smilek, Ristic, Friesen, & Eastwood, 2003; Papeo, Stein, & Soto-Faraco, 2017) as important in determining some aspects of visual scene perception. These factors are to play an especially important role in real-life naturalistic scenarios, where these high-level relationships are often abundant (Peelen & Kastner, 2014). Adding to this evidence from visual-only experiments, in the present study we demonstrated that high-level cross-modal (auditory-visual) semantic relations may as well exert an impact in spatial orienting and guide attention in visual search for objects in real-life, dynamic scenes. In fact, one could speculate that especially in complex and noisy environments where many visual and auditory events are spatially and temporally coincident, semantic information might become a leading predictor of object presence, and hence, guide attention.

One open question is why target-consistent sounds benefit search, but distracter-consistent sounds do not slow down reaction times (in comparison to neutral or no sounds). If cross-modal interactions were strictly automatic and pre-attentive, then distractor sounds should increase the saliency of their corresponding, yet irrelevant objects present in the scene. However, the evidence we found is not consistent with the strong pre-attentive view of cross-modal semantic effects. Despite the interplay between attention and multisensory interactions is far from resolved (Talsma et al., 2010; ten Oever et al., 2016; Hartcher-O’Brien et al., 2017; Lunn et al., 2019; Soto-Faraco et al., in press, for some reviews), many studies illustrate the multisensory interactions tend to wane when the implicated inputs are not attended (e.g., Alsius et al., 2005; 2014; Talsma & Woldorff, 2006). For example, Molholm et al., (2004) demonstrated that object-based enhancement occurs in a goal-directed manner, suggesting that while a characteristic sound of a target will facilitate its localization, a characteristic sound of a distracter will not attract attention to the distracter. Consistent with the idea of automaticity (and therefore, contrary Molholm et al. and to our results), a study by Mastroberardino et al. (2015) showed that audio-visual events can capture attention even when not task-relevant. One main difference between studies is that Mastroberardino et al. used a low perceptual load situation with a very limited range of possible semantic relationships (just two). Therefore, object-based enhancements might eventually occur even when task irrelevant, under favourable low load conditions. Further studies to understand the limits of crossmodal semantic effects and how they apply to real-life dynamic scenarios should be ran to clarify this point. For example, in line with the present study, a possible next step would be to use eye tracking with free-viewing of the video-clips to investigate if crossmodal semantic congruency attracts visual behaviour and can be, therefore, responsible for the visual search effects seen here.

In conclusion, we have demonstrated that semantic consistent sounds can produce an enhancement in visual search in complex and dynamic scenes. We suggest that this enhancement happens through object-based interactions between visual and auditory modalities. This demonstration not only generalizes (and confirms) previous laboratory findings on semantically-based crossmodal interactions but expands it to the field of research in natural scenes.

## Supporting information

example sounds

Example video 1

Example video 1

## Acknowledgements

This research was supported by the Ministerio de Economia y Competitividad (PSI2016-75558-P AEI/FEDER), AGAUR Generalitat de Catalunya (2017 SGR 1545). Daria Kvasova was supported by an FI scholarship, from the AGAUR Generalitat de Catalunya.

## References

Biederman, I., Mezzanotte, R. J., and Rabinowitz, J. C. (1982). Scene perception: detecting and judging objects undergoing relational violations. Cogn. Psychol. 14, 143–177.

Bolognini, N., Frassinetti, F., Serino, A., & Làdavas, E. (2005). “Acoustical vision” of below threshold stimuli: Interaction among spatially converging audiovisual inputs. Experimental Brain Research, 160(3), 273–282. https://doi.org/10.1007/s00221-004-2005-z

Busse, L., Roberts, K. C., Crist, R. E., Weissman, D. H., & Woldorff, M. G. (2005). The spread of attention across modalities and space in a multisensory object. Proceedings of the National Academy of Sciences, 102(51), 18751–18756. https://doi.org/10.1073/pnas.0507704102

Chen, Y., & Spence, C. (2011). Crossmodal Semantic Priming by Naturalistic Sounds and Spoken Words Enhances Visual Sensitivity. Journal of Experimental Psychology: Human Perception and Performance, 37(5), 1554–1568. https://doi.org/10.1037/a0024329

Chun, M. M. (2000). Contextual cueing of visual attention. Trends in Cognitive Sciences, 4(5), 170–178. https://doi.org/10.1016/S1364-6613(00)01476-5

Driver, J., & Spence, C. (2011). Cross-modal links in spatial attention of Cognitive. Society, 353(1373), 1319–1331. https://doi.org/10.1098/rstb.1998.0286

Evans, K.K., Georgian-Smith D., Tambouret R., Birdwell R.L., & Wolfe J.M. (2013) The gist of the abnormal: above-chance medical decision making in the blink of an eye. Psychonomic Bulletin & Review, 20, 1170–1175. https://doi.org/10.3758/s13423-013-0459-3

Frassinetti, F., Bolognini, N., & Làdavas, E. (2002). Enhancement of visual perception by crossmodal visuo-auditory interaction. Experimental Brain Research, 147(3), 332–343. https://doi.org/10.1007/s00221-002-1262-y

Green, C., Hummel, J. E. (2004). Relational perception and cognition: Implications for cognitive architecture and the perceptual-cognitive interface. In Ross, B. H.(Ed.), The psychology of learning and motivation (Vol. 44, pp. 201–226).

Green, C., Hummel, J. E. (2006). Familiar interacting object pairs are perceptually grouped. Journal of Experimental Psychology: Human Perception and Performance, 32, 1107–1119. http://dx.doi.org/10.1037/0096-1523.32.5.1107

Greene, M.R. and Oliva, A. (2009) Recognition of natural scenes from global properties: seeing the forest without representing the trees. Cogn. Psychol. 58, 137–176. https://doi/10.1037/0096-1523.32.5.110

Hartcher-O’Brien, J., Soto-Faraco, S., & Adam, R. (2017). a matter of bottom-up or top-down processes: the role of attention in multisensory integration. Frontiers in integrative neuroscience, 11, 5. https://doi/10.3389/fnint.2017.00005

Hasson, U., Malach, R., & Heeger, D. J. (2010). Reliability of cortical activity during natural stimulation. Trends in Cognitive Sciences, 14(1), 40–48. https://doi/10.1016/j.tics.2009.10.011

Hershler, O., & Hochstein, S. (2009). The importance of being expert: Top-down attentional control in visual search with photographs. Attention, Perception, & Psychophysics, 71(7), 1478–1486. https://doi/10.3758/APP.71.7.1478

Iordanescu, L., Guzman-Martinez, E., Grabowecky, M., & Suzuki, S. (2008). Characteristic sounds facilitate visual search. Psychonomic Bulletin & Review, 15(3), 548–54. https://doi.org/10.3758/PBR.15.3.548

Iordanescu, L., Grabowecky, M., Franconeri, S., Theeuwes, J., & Suzuki, S. (2010). Characteristic sounds make you look at target objects more quickly. Attention, Perception, & Psychophysics, 72(7), 1736–1741. https://doi/10.3758/APP.72.7.1736

Kaiser, D., Stein, T., & Peelen, M. V. (2014). Object grouping based on real-world regularities facilitates perception by reducing competitive interactions in visual cortex. Proceedings of the National Academy of Sciences, 111(30), 11217–11222. https://doi/10.1073/pnas.1400559111

Kingstone, A., Smilek, D., Ristic, J., Friesen, C. K., & Eastwood, J. D. (2003). Attention, Researchers! It Is Time to Take a Look at the Real World. Current Directions in Psychological Science, 12(5), 176–180. https://doi.org/10.1111/1467-8721.01255

Knoeferle, K. M., Knoeferle P., Velasco C., & Spence C. (2016). Multisensory brand search: how the meaning of sounds guides consumers’ visual attention. Journal of Experimental Psychology. 22(2):196–210. https://doi.org/10.1037/xap0000084

Kuai, S.G., Levi D., & Kourtzi Z. (2013) Learning optimizes decision templates in the human visual cortex. Current Biology, 23, 1799–1804. https://doi.org/10.1016/j.cub.2013.07.052

List, A., Iordanescu, L., Grabowecky, M., & Suzuki, S. (2014). Haptic guidance of overt visual attention. Attention, Perception, & Psychophysics, 76(8), 2221–2228. https://doi.org/10.3758/s13414-014-0696-1

Lunn, J., Sjoblom, A., Ward, J., Soto-Faraco, S., & Forster, S. (2019). Multisensory enhancement of attention depends on whether you are already paying attention. Cognition, 187, 38–49. https://doi.org/10.1016/j.cognition.2019.02.008

Maddox, R. K., Atilgan, H., Bizley, J. K., & Lee, A. K. (2015). Auditory selective attention is enhanced by a task-irrelevant temporally coherent visual stimulus in human listeners. Elife, 4, e04995. https://doi.org/10.7554/eLife.04995

Maguire, E. A. (2012). Studying the freely-behaving brain with fMRI. Neuroimage, 62(2), 1170–1176. https://doi.org/10.1016/j.neuroimage.2012.01.009

Malinowski, P., & Hübner, R. (2001). The effect of familiarity on visual-search performance: Evidence for learned basic features. Attention, Perception, & Psychophysics, 63(3), 458–463. https://doi.org/10.3758/BF03194412

Mastroberardino, S., Santangelo, V., & Macaluso, E. (2015). Crossmodal semantic congruence can affect visuo-spatial processing and activity of the fronto-parietal attention networks. Frontiers in Integrative Neuroscience, 9(July), 45. https://doi.org/10.3389/fnint.2015.00045

Matusz, P.J., Dikker S., Huth A.G & Perrodin C. (2019). Are we ready for real-world neuroscience? Journal of Cognitive Neuroscience, 31(3), 327–338. https://doi.org/10.1162/10.1162/jocn_e_01276

McDonald, J. J., Teder-Salejarvi, W. A., & Hillyard, S. A. (2000). Involuntary orienting to sound improves visual perception. Nature, 407(6806), 906. https://doi.org/10.1038/35038085

Molholm, S., Ritter, W., Javitt, D. C., & Foxe, J. J. (2004). Multisensory Visual-Auditory Object Recognition in Humans: A High-density Electrical Mapping Study. Cerebral Cortex, 14(4), 452–465. https://doi.org/10.1093/cercor/bhh007

Nardo, D., Santangelo, V., and Macaluso, E. (2014). Spatial orienting in complex audiovisual environments. Human brain mapping, 35, 1597–1614. https://doi.org/10.1002/hbm.22276

Neisser, U. (1976). Cognition and reality: Principles and implications of cognitive psychology. WH Freeman/Times Books/Henry Holt & Co. https://doi.org/10.1068/p060605

Nickerson, R. S. (1973). Intersensory facilitation of reaction time: Energy summation or preparation enhancement? Psychological Review, 80 489–50.

Olivers, C. N. L. and van der Burg, E. (2008). Bleeping you out of the blink: sound saves vision from oblivion, Brain Research, 1242, 191–199. http://dx.doi.org/10.1016/j.brainres.2008.01.070.

Oliva, A. and Torralba, A. (2007) The role of context in object recognition. Trends Cogn. Sci. 11, 520–527. https://doi.org/10.1016/j.tics.2007.09.009

Papeo L., Stein T., Soto-Faraco S. (2017). The two-body inversion effect. Psychological Science, 28(3), 369–379. https://doi.org/10.1177/0956797616685769

Peelen, M. V., Fei-Fei, L., & Kastner, S. (2009). Neural mechanisms of rapid natural scene categorization in human visual cortex. Nature, 460(7251), 94. https://doi.org/10.1038/nature08103

Peelen, M. V., & Kastner, S. (2014). Attention in the real world: Toward understanding its neural basis. Trends in Cognitive Sciences, 18(5), 242–250. https://doi.org/10.1016/j.tics.2014.02.004

Perazzo, E. (2017). Iron Fist, New York, USA.: Marvel Television in association with ABC Studios.

Pesquita, A., Brennan, A. A., Enns, J. T., & Soto-Faraco, S. (2013). Isolating shape from semantics in haptic-visual priming. Experimental Brain Research, 227(3), 311–322. http://dx.doi.org/10.1007/s00221-013-3489-1

Purves, D., Wojtach, W. T., & Lotto, R. B. (2011). Understanding vision in wholly empirical terms. Proceedings of the National Academy of Sciences, 108(Supplement 3), 15588–15595. https://doi.org/10.1073/pnas.1012178108

Ramos-Estebanez, C., Merabet, L. B., Machii, K., Fregni, F., Thut, G., Wagner, T. A., & Pascual-Leone, A. (2007). Visual phosphene perception modulated by subthreshold crossmodal sensory stimulation. Journal of Neuroscience, 27(15), 4178–4181. https://doi.org/10.1523/JNEUROSCI.5468-06.2007

Shiffrin, R. M., & Schneider, W. (1977). Controlled and automatic human information processing: II. Perceptual learning, automatic attending and a general theory. Psychological Review, 84(2), 127. http://dx.doi.org/10.1037/0033-295X.84.2.12

Soto-Faraco, S., Biau, E., Moris-Fernandez, L., Ikumi, N., Kvasova, D., Ruzzoli, M. & Torralba, M. (in press). Multisensory integration in the real world. To appear in M. Chun(Ed.), Cambridge Elements of Perception. Cambridge, UK: Cambridge University Press.

Spence, C. J., & Driver, J. (1994). Covert spatial orienting in audition: Exogenous and endogenous mechanisms. Journal of Experimental Psychology: Human Perception and Performance, 20(3), 555–574. http://dx.doi.org/10.1037/0096-1523.20.3.555

Spence, C., and Driver, J. (2004). Crossmodal Space and Crossmodal Attention (Oxford: Oxford University Press). https://doi.org/10.1093/acprof:oso/9780198524861.001.0001

Stein, B. E., & Meredith, M. A. (1993). The merging of the senses. The MIT Press. https://doi.org/10.1177/027836499401300508

Talsma, D., Senkowski, D., Soto-Faraco, S., & Woldorff, M. G. (2010). The multifaceted interplay between attention and multisensory integration. Trends in cognitive sciences, 14(9), 400–410. https://doi.org/10.1016/j.tics.2010.06.008

Teder-Sälejärvi, W. A., Di Russo, F., McDonald, J. J., & Hillyard, S. A. (2005). Effects of spatial congruity on audio-visual multimodal integration. Journal of cognitive neuroscience, 17(9), 1396–1409. https://doi.org/10.1162/0898929054985383

Ten Oever, S., Romei, V., van Atteveldt, N., Soto-Faraco, S., Murray, M. M., & Matusz, P. J. (2016). The COGs (context, object, and goals) in multisensory processing. Experimental brain research, 234(5), 1307–1323. https://dx.doi.org/10.1007/s00221-016-4590-z

van den Brink, R. L.,Cohen, M.X., van der Burg, E., Talsma, D., Vissers, M.E., and Slagter, H.A. (2014). Subcortical, modality-specific pathways contribute to multisensory processing in humans. Cereb. Cortex 24, 2169–2177. https://doi.org/10.1093/cercor/bht069

Van der Burg, E., Olivers, C. N. L., Bronkhorst, A. W., & Theeuwes, J. (2008). Pip and pop: Nonspatial auditory signals improve spatial visual search. Journal of Experimental Psychology-Human Perception and Performance, 34(5), 1053–1065. https://doi.org/10.1037//0096-1523.26.5.1583

Vatakis, Argiro and Charles Spence (2010). Audiovisual Temporal Integration for Complex Speech, Object-Action, Animal Call, and Musical Stimuli in Multisensory Object Perception in the Primate Brain, ed. M. J. Naumer and J. Kaiser: Springer, 95–121 https://doi.org/10.3389/fnint.2012.00071

Vroomen, J. and de Gelder, B. (2000). Sound enhances visual perception: crossmodal effects of auditory organization on vision, Journal of Experimental Psychology: Human Perception and Performance, 26, 1583–1590. https://doi.org/10.1037//0096-1523.26.5.1583

Wang, Q., Cavanagh, P., & Green, M. (1994). Familiarity and pop-out in visual search. Attention, Perception, & Psychophysics, 56(5), 495–500. https://doi.org/10.1037//0096-1523.26.5.1583

Wolfe, J. M., Horowitz, T. S., & Kenner, N. M. (2005). Cognitive psychology: rare items often missed in visual searches. Nature, 435(7041), 439. https://doi.org/10.1038//435439a

Wu, C., Wick, F. A., & Pomplun, M. (2014). Guidance of visual attention by semantic information in real-world scenes. Frontiers in Psychology, 5(February), 1–13. https://doi.org/10.3389/fpsyg.2014.00054

